# Cryptic β-lactamase evolution is driven by low β-lactam concentrations

**DOI:** 10.1101/2020.12.01.404343

**Authors:** Christopher Fröhlich, João Alves Gama, Klaus Harms, Viivi H.A. Hirvonen, Bjarte Aarmo Lund, Marc W. van der Kamp, Pål Jarle Johnsen, Ørjan Samuelsen, Hanna-Kirsti S. Leiros

**Affiliations:** The Norwegian Structural Biology Centre (NorStruct), Department of Chemistry, UiT The Arctic University of Norway, Tromsø, Norway; Department of Pharmacy, UiT The Arctic University of Norway, Tromsø, Norway; School of Biochemistry, University of Bristol, Bristol, UK; Norwegian National Advisory Unit on Detection of Antimicrobial Resistance, Department of Microbiology and Infection Control, University Hospital of North Norway, Tromsø, Norway

**Author notes:** These authors contributed equally (Ørjan Samuelsen, Hanna-Kirsti S. Leiros).

**Keywords:** OXA-48, ceftazidime, resistance development, cryptic evolution, *Escherichia coli*, carbapenemase, carbapenem, sub-MIC, structural flexibility, catalytic efficiency

## Abstract

Our current understanding of how low antibiotic concentrations shape the evolution of contemporary β-lactamases is limited. Using the wide-spread carbapenemase OXA-48, we tested the long-standing hypothesis that selective compartments with low antibiotic concentrations cause standing genetic diversity that could act as a gateway to develop clinical resistance. Here, we subjected *Escherichia coli* expressing *bla*_OXA-48_, on a clinical plasmid, to experimental evolution at sub-minimum inhibitory concentrations (sub-MIC) of ceftazidime. We identified and characterized seven single variants of OXA-48. Susceptibility profiles and dose-response curves showed that they increased resistance only marginally. However, in competition experiments at sub-MIC of ceftazidime, they showed strong selectable fitness benefits. Increased resistance was also reflected in elevated catalytic efficiencies towards ceftazidime. These changes are likely caused by enhanced flexibility of the Ω- and β5-β6 loops. In conclusion, low-level concentrations of β-lactams can drive the evolution of β-lactamases through cryptic phenotypes which may act as stepping-stones towards clinical resistance.

## INTRODUCTION

Since the discovery of the first β-lactam, penicillin, this antibiotic class has diversified into a broad range of agents and it remains the most widely used class of antibiotics worldwide (Bush & Bradford, 2016). The extensive use of these agents has inevitably led to the selection of multiple resistance mechanisms where the expression of β-lactamase enzymes plays a major role, particularly in Gram-negative bacteria (Bush, 2018). Consequently, β-lactamases are arguably among the most studied enzymes world-wide. Considerable progress has been made in understanding their molecular epidemiology and biochemical properties (Bonomo, 2017; Pitout et al., 2019). The evolutionary forces driving the diversification of these enzymes are however poorly understood. Already more than twenty years ago, it was proposed that sub-optimal antibiotic concentrations within the host fuel the evolution of β-lactamases, altering their substrate profiles (Baquero, 2001; Baquero & Negri, 1997; Baquero et al., 1997; Negri et al., 2000). This “compartment hypothesis” was later supported by a series of studies unequivocally demonstrating that selection for antibiotic resistance determinants can occur at very low antibiotic concentrations (Gullberg et al., 2014; Gullberg et al., 2011; Westhoff et al., 2017). Despite their clinical significance, few studies have investigated the effects of sub-minimum inhibitory concentrations (sub-MIC) of β-lactams on the evolution and selection of contemporary, globally circulating β-lactamases (Bagge et al., 2004; Murray et al., 2018; Negri et al., 2000).

Within the last decade, OXA-48 has become one of the most widespread serine β-lactamases. This Ambler class D β-lactamase confers resistance towards penicillins and decreases susceptibility to our last-resort drugs, the carbapenems. However, it is ineffective against extended-spectrum cephalosporins including ceftazidime (Docquier et al., 2009; Fröhlich et al., 2019; Poirel et al., 2004). Despite that, naturally occurring OXA-48-like variants have been identified exhibiting increased ceftazidime activity but limited hydrolytic activity towards penicillins and carbapenems (e.g. OXA-163, OXA-247 and OXA-405) (Dortet et al., 2015; Gomez et al., 2013; Poirel et al., 2011). Ceftazidime resistance development in these variants was mostly due to single amino acid changes and a shortened β5-β6 loop (Dortet et al., 2015; Gomez et al., 2013; Mairi et al., 2018; Pitout et al., 2019; Poirel et al., 2011). We previously showed that exposure to increasing concentrations of ceftazidime can select for this latent ceftazidimase function of OXA-48 in the laboratory (Fröhlich et al., 2019).

To test the long-standing hypothesis, that β-lactams at sub-MIC can drive the evolution of these enzymes, we subjected *Escherichia coli* MG1655 expressing OXA-48 to concentrations of ceftazidime below the MIC (0.25xMIC). Over the course of 300 generations, we identified seven single variants of OXA-48 (L67F, P68S, F72L, F156C/V, L158P and G160C). Their ceftazidime MIC were indistinguishable or only marginally increased, compared to wild-type OXA-48. However, when expressed at sub-MIC of ceftazidime, all allele variants conferred strong fitness benefits. Measuring dose-response curves (IC_50_) and enzyme kinetics revealed further that (i) all genotypes decreased ceftazidime susceptibility significantly and (ii) all enzyme variants exhibited increased catalytic efficiencies against ceftazidime. Molecular dynamics (MD) simulations of P68S, F72L and L158P showed elevated flexibility of both the Ω- (D143 to I164) and β5-β6 (T213 to K218) loops likely to aid hydrolysis of the bulkier ceftazidime by increasing active site accessibility. Structural investigations of L67F also revealed a novel binding site for the hydrolysed ceftazidime where the β5-β6 loop was also involved in the product release. Worryingly, double mutants, such as F72I/G131 (OXA-D320, GenBank accession no. KJ620465) and N146S/L158P (OXA-D319, GenBank accession no. KJ620462), were recently identified in environmental samples (Naas et al., 2017; Tacao et al., 2017) underlining the importance and evolutionary power of environments with low-selective pressure.

## RESULTS

### Sub-MIC of ceftazidime select for high-level resistance

Here, we wanted to study the evolvability of the carbapenemase OXA-48 under sub-MIC of the cephalosporin ceftazidime. OXA-48 does not hydrolyse ceftazidime efficiently (Poirel et al., 2011). However, we recently showed, that the exposure to increasing concentrations of ceftazidime can select for OXA-48 variants with elevated activity towards ceftazidime (Fröhlich et al., 2019). We used the previously constructed *E. coli* MG1655 (Table S1, MP13-06) (Fröhlich et al., 2019) carrying a globally disseminated IncL plasmid with *bla*_OXA-48_ as the only antibiotic resistance gene. MP13-06 was evolved without selection pressure and at one quarter of the ceftazidime MIC (0.06 mg/L) resulting in the populations 1 to 3 (Pop1 to 3) and 4 to 6 (Pop4 to 6), respectively (Figure 1A).

**Figure 1.**
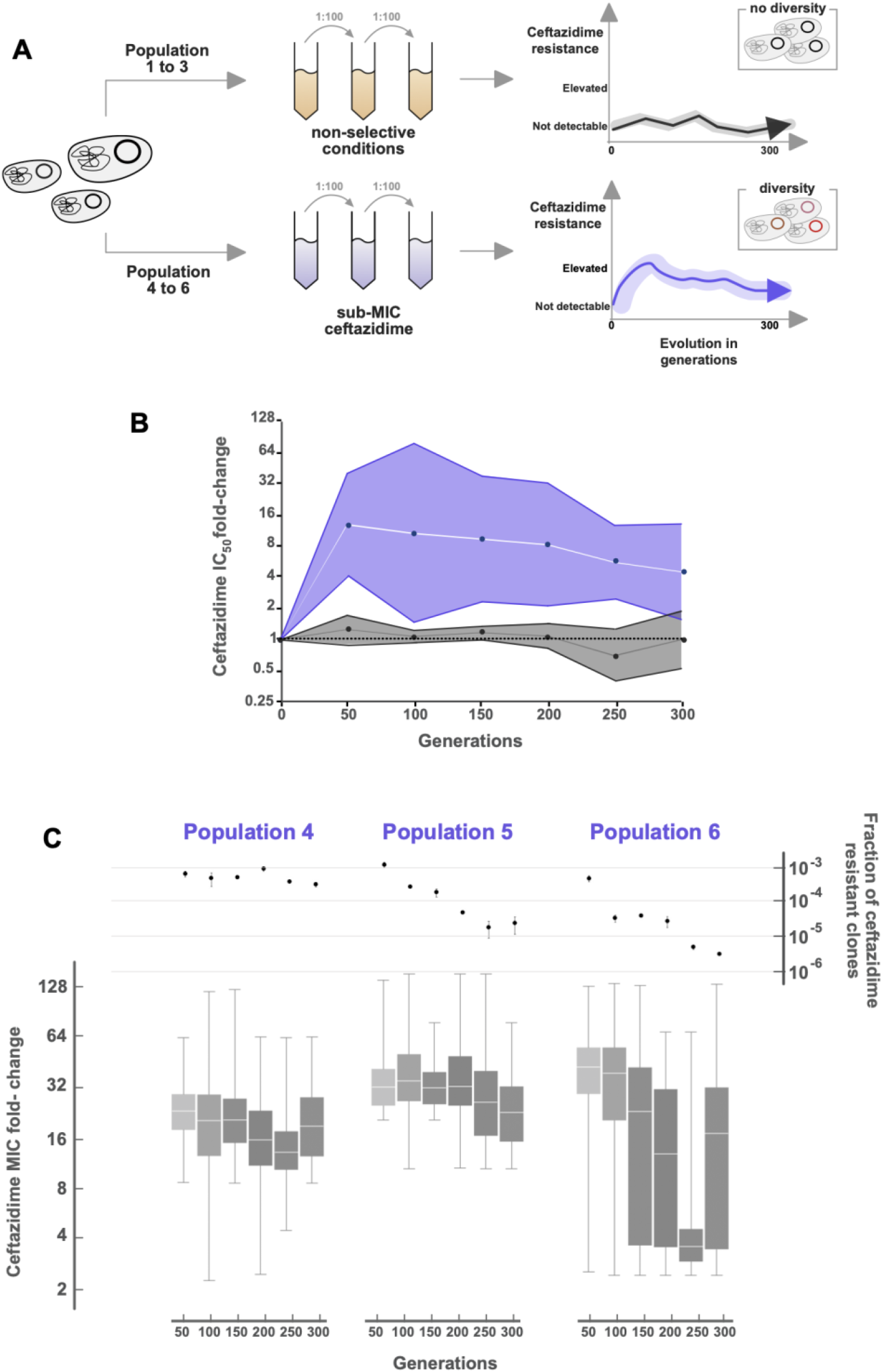
Population level effects of sub-MIC ceftazidime exposure. A. Experimental design. B: IC_50_ fold-change for populations evolved without (grey) and under sub-MIC ceftazidime conditions (violet), relative to wild-type OXA-48. Bands represent the standard deviation around the geometric mean. C: Top section shows the fraction of clones able to grow on ceftazidime 1 mg/L (>2-fold MIC). Bottom section displays MIC fold-change distributions of pre-selected clones. Boxplots represent quartiles and the median of the distributions.

To elucidate the effect of ceftazidime, we measured dose-response curves of the whole evolved populations and calculated the ceftazidime concentrations inhibiting 50% of cell growth (IC_50_). In Pop1 to 3, evolution without selection pressure did not result in altered ceftazidime susceptibility (Figure 1B). In contrast, under ceftazidime selection (Pop4 to 6), susceptibility decreased on average 16-fold already after 50 generations (Figure 1B). We observed that, during the course of experimental evolution, the susceptibilities of Pop4 to 6 shifted towards lower ceftazidime resistance (Figure 1B).

From the evolved populations, we measured the fraction of clones exhibiting a clinically significant MIC change (>2-fold) by non-selective and selective plating on 1 mg/L ceftazidime. No clones were identified during selection-free evolution above the detection limit (10^-7^ of the population). Under sub-MIC conditions, we found a significant fraction of the populations able to grow on ceftazidime containing plates (Figure 1C). While this fraction was stably maintained in Pop4, we found that Pop5 and Pop6 showed a significant reduction over time (Pearson correlation, P=0.54, P=0.01, P=0.03).

To determine the MIC distribution of clones with increased MIC, we selected approximately 50 colonies every 50^th^ generation from the selective plates and tested their susceptibility to ceftazidime (Figure 1C). All pre-selected clones displayed a MIC increase ranging from 2- to 128-fold. For Pop 4 to 6, we found that on average 11%, 28% and 34% of the tested colonies exhibited MIC values above the clinical resistance breakpoint of 4 mg/L, respectively (Breakpoint table v. 10.0). These results are consistent with recent reports demonstrating that low-level concentrations of antibiotics facilitate the selection of high-level resistance (Gullberg et al., 2011; Westhoff et al., 2017).

### Sub-MIC evolution selects for beneficial single point mutations in *bla*_OXA-48_

To understand the effect of sub-MIC exposure on OXA-48, we sequenced the *bla*_OXA-48_ gene of approximately 50 clones after 50 and 300 generations, which were pre-selected on agar plates containing 1 mg/L ceftazidime. In total, seven single variants of OXA-48 were identified: L67F, P68S, F72L, F156C, F156V, L158P and G160C. The relative frequency of these variants varied among populations and generations (Figure 2A). Interestingly, double mutants with similar (F72I/G131S) or identical (N146S/L158P) amino acid changes have been already reported in environmental samples (Tacao et al., 2017). To elucidate the effect of these second mutations, we constructed the OXA-48 double mutants F72L/G131S (instead of F72I) and N146S/L158P and included them in the following characterization.

**Figure 2.**
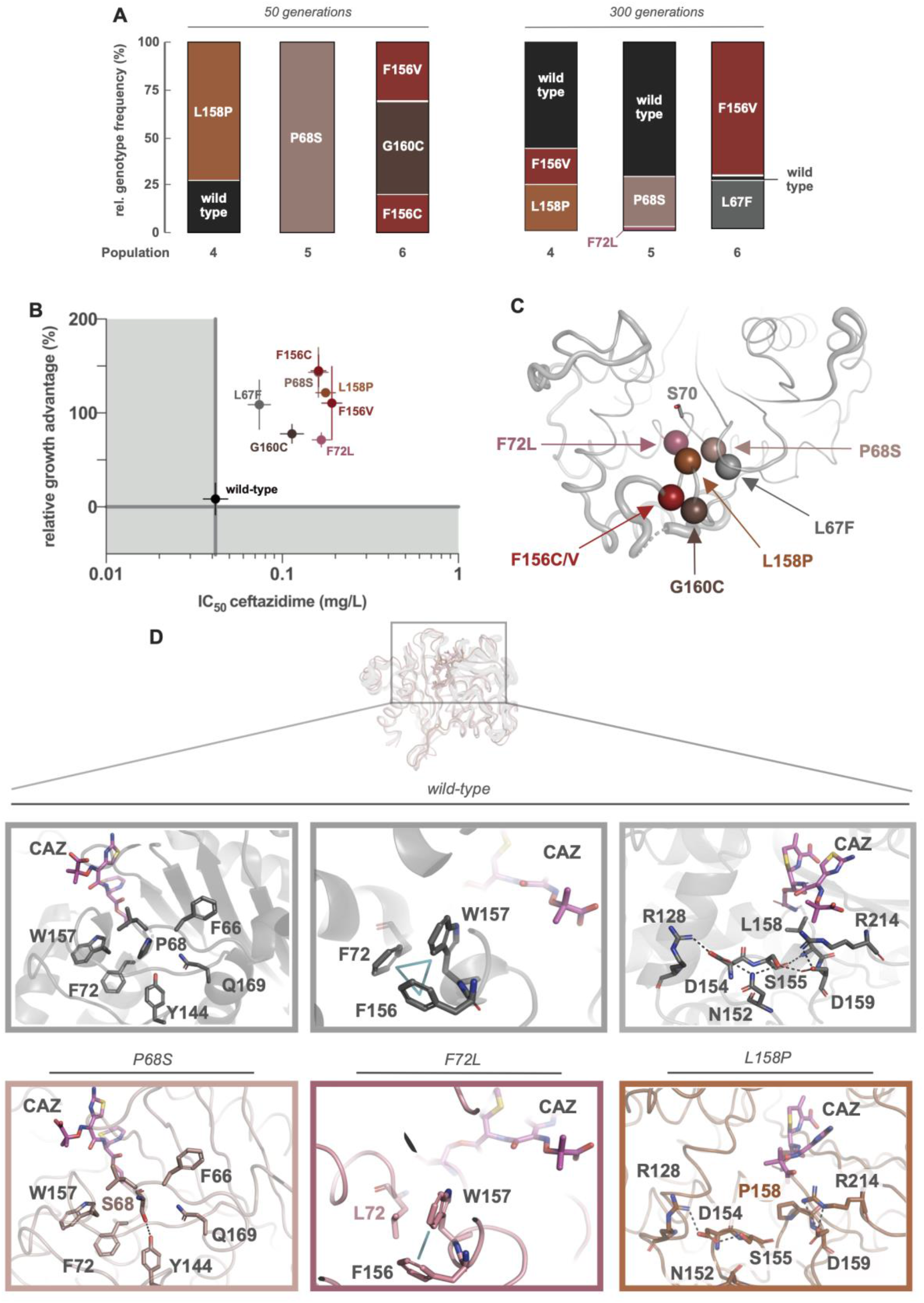
Phenotypic and structural investigation of OXA-48 allele variants. A. Relative genotype frequencies of *bla*_OXA-48_ variants within Pop4 to 6. B. Relative growth advantage of OXA-48 variants expressed at sub-MIC of ceftazidime *versus* their ceftazidime IC_50_. Despite marginal changes in their ceftazidime susceptibility (IC_50_ increased by 2- to 4-fold), the expression of these alleles displays large fitness benefits at sub-MIC ceftazidime. Error bars represent the standard deviation. C. Ribbon structure of OXA-48 including the amino acid changes close to the active site. D. Representative structures from molecular dynamics simulations of wild-type, P68S, F72L and L158P performed with ceftazidime covalently bound to the active site S70. In short, S68 in P68S displays an H-bond with the tyrosine in the conserved Y^144^GN motif of OXA-48. F72L lacks the aromatic stacking interaction between F72 and F156/W157. L158P disrupts the H-bond network within the Ω-loop.

To isolate the effects of OXA-48 on antimicrobial susceptibility, we sub-cloned all allele variants into a high-copy number vector (pCR-Blunt II-TOPO) and expressed them in *E. coli* TOP10 (Table 1). As previously shown (Fröhlich et al., 2019), the expression of OXA-48 resulted in up to 32- and 64-fold increased MIC towards penicillins and carbapenems (except for doripenem), respectively. While wild-type OXA-48 expression did not increase the MIC against cephalosporins (<2-fold), we found that P68S, F72L, L158P and N146S/L158P resulted in 4- to 16-fold increased MIC against ceftazidime. Interestingly, the expression of all other alleles (L67F, F156C/V, G160C, F72L/G131S) did not increase the ceftazidime MIC significantly (i.e., not more than 2-fold, compared to wild-type OXA-48). In addition, none of the alleles showed a significant effect on cephalosporins other than ceftazidime (Table 1). In contrast, the susceptibility to carbapenems and penicillins was increased by 2- to ≥64-fold for all the variants. The expression of F156C and G160C did not increase the MIC to any β-lactam. In clinical strains, OXA-48 is frequently located on IncL plasmids which are typically present in low copy numbers (Preston et al., 2014). To mimic this situation, we sub-cloned all OXA-48 alleles into a low copy number vector (pUN), expressed these in *E. coli* MG1655, and repeated the ceftazidime MIC measurements. Within this more realistic genetic architecture only F156V increased the ceftazidime MIC by more than 2-fold (Table S2).

**Table 1.** MIC (mg/L) of OXA-48 and allele variants expressed in the high copy number vector pCR-Blunt II-TOPO in *E. coli* TOP10

| Antimicrobial agents<sup>a</sup> | MP13-04 | MP13-11<br>wild-type<br>OXA-48 | MP13-21<br>L67F | MP13-16<br>P68S | MP13-14<br>F72L | MP13-17<br>F156C | MP13-18<br>F156V | MP13-15<br>L158P | MP13-19<br>G160C | MP13-33<br>F72L/<br>G131S | MP13-20<br>N146S/<br>L158P |
| --- | --- | --- | --- | --- | --- | --- | --- | --- | --- | --- | --- |
| Temocillin | 16 | 256 | 64 | 64 | 64 | 16 | 16 | 64 | 16 | 16 | 32 |
| Piperacillin/tazobactam | 2 | 64 | 2 | 2 | 2 | 2 | 2 | 2 | 2 | 2 | 2 |
| Amoxicillin/clavulanic acid | 4 | 128 | 128 | 64 | 64 | 16 | 64 | 16 | 8 | 8 | 8 |
| Ceftazidime | 0.5 | 0.5 | 1 | 8 | 4 | 0.5 | 0.5 | 8 | 0.5 | 0.5 | 2 |
| Ceftazidime/avibactam | 0.25 | 0.25 | 0.5 | 0.25 | 0.5 | 0.5 | 0.5 | 0.5 | 0.5 | 0.25 | 0.5 |
| Cefuroxime | 16 | 16 | 8 | 16 | 8 | 8 | 8 | 8 | 8 | 16 | 8 |
| Cefepime | 0.06 | 0.12 | 0.06 | 0.12 | 0.12 | 0.06 | 0.06 | 0.25 | 0.06 | 0.06 | 0.12 |
| Cefotaxime | 0.12 | 0.25 | 0.12 | 0.12 | 0.12 | 0.12 | 0.12 | 0.25 | 0.12 | 0.25 | 0.12 |
| Meropenem | 0.03 | 0.25 | 0.12 | 0.03 | 0.03 | 0.03 | 0.03 | 0.03 | 0.03 | 0.03 | 0.03 |
| Imipenem | 0.25 | 1 | 0.25 | 0.25 | 0.25 | 0.25 | 0.25 | 0.25 | 0.25 | 0.25 | 0.5 |
| Ertapenem | 0.015 | 1 | 0.12 | 0.03 | 0.03 | 0.015 | 0.015 | 0.06 | 0.015 | 0.015 | 0.015 |
| Doripenem | 0.03 | 0.03 | 0.03 | 0.03 | 0.06 | 0.03 | 0.03 | 0.06 | 0.03 | 0.06 | 0.06 |

<sup>a</sup> tazobactam fixed at 4 µg/mL; clavulanic acid fixed at 2 µg/mL; avibactam fixed at 4 µg/mL

Clinically insignificant or marginal changes in MIC have been reported to still confer high fitness benefits in the presence of low-level selection (Negri et al., 2000). To address this, we first increased the resolution of the susceptibility testing by measuring the dose-response curves, in the low copy number vector. Calculating their corresponding IC50 values, we found that all variants conferred marginal but significant decreases in ceftazidime susceptibility (Figure 2B and Table S2, ANOVA, df=10, P<0.0001, followed by Dunnett post hoc test).

Secondly, we performed head-to-head competitions between isogenic *E. coli* MG1655 strains (only differing in Δ*malF*) to test the fitness effect of OXA-48 variants in the absence and presence of sub-MIC ceftazidime. To exclude an effect of the *malF* deletion on the bacterial fitness, we initially competed the strains both carrying the pUN vector encoding wild-type *bla*_OXA-48_ (MP08-61 and MP14-24). No significant change in bacterial fitness was observed in either condition (Welch t-test, P=0.24 and P=0.48), out-ruling a detectable effect of the *malF* deletion. Therefore, we next expressed all OXA-48 variants in MP14-23 and subjected those to competitions against MP08-61. Without ceftazidime, no difference in fitness was observed between variants and wild-type (Figure S1; ANOVA, not assuming equal variances, df=7, P=0.33). However, at sub-MIC ceftazidime, all allele variants showed strong significant growth benefits (Figure 2B; ANOVA, not assuming equal variances, df=7, P=0.0003, followed by a Dunnett post hoc test with OXA-48 as control group).

Two mutational targets identified in our study (F72L and L158P) were recently isolated from the environment, in combination with a second amino acid substitution (Tacao et al., 2017). We aimed to elucidate the effect of G131S and N146S in combination with F72L and L158P, respectively. To do so, we competed the F72L and L158P against the double mutants F72L/G131S and N146S/L158P, respectively. No significant change in fitness was detectable in the absence of selection pressure (Figure S2; Welch t-test, P=0.24 and 0.62 for F72L/G131S and N146S/L158P, respectively). At sub-MIC ceftazidime, our data suggest no positive selection for the double mutants (Figure S2; paired Welch t-test between conditions, P=0.038 and 0.59 for F72L/G131S and N146S/L158P). Thus, the role of these second mutations remains unclear.

### Mutations within *bla*_OXA-48_ alter enzymatic properties

In this study, sub-MIC ceftazidime were shown to select for OXA-48 variants conferring high bacterial fitness advantages despite only cryptic resistance phenotypes suggesting that, in clinic set-ups, these genetic changes are likely to remain undetected. To further our understanding of how these cryptic changes influence the enzymatic properties of OXA-48, we expressed the enzymes without their leader sequence in *E. coli* BL21 AI (Table S1). After enzyme purification, protein masses were verified using electrospray ionization mass spectrometry (ESI-MS). The molecular weight of five out of seven variants corresponded to their calculated monoisotopic masses (Table 2). For F156C and G160C, we observed an increase in molecular weight by 76 Da, likely caused by β-mercaptoethanol used during the purification process to increase solubility.

**Table 2.**
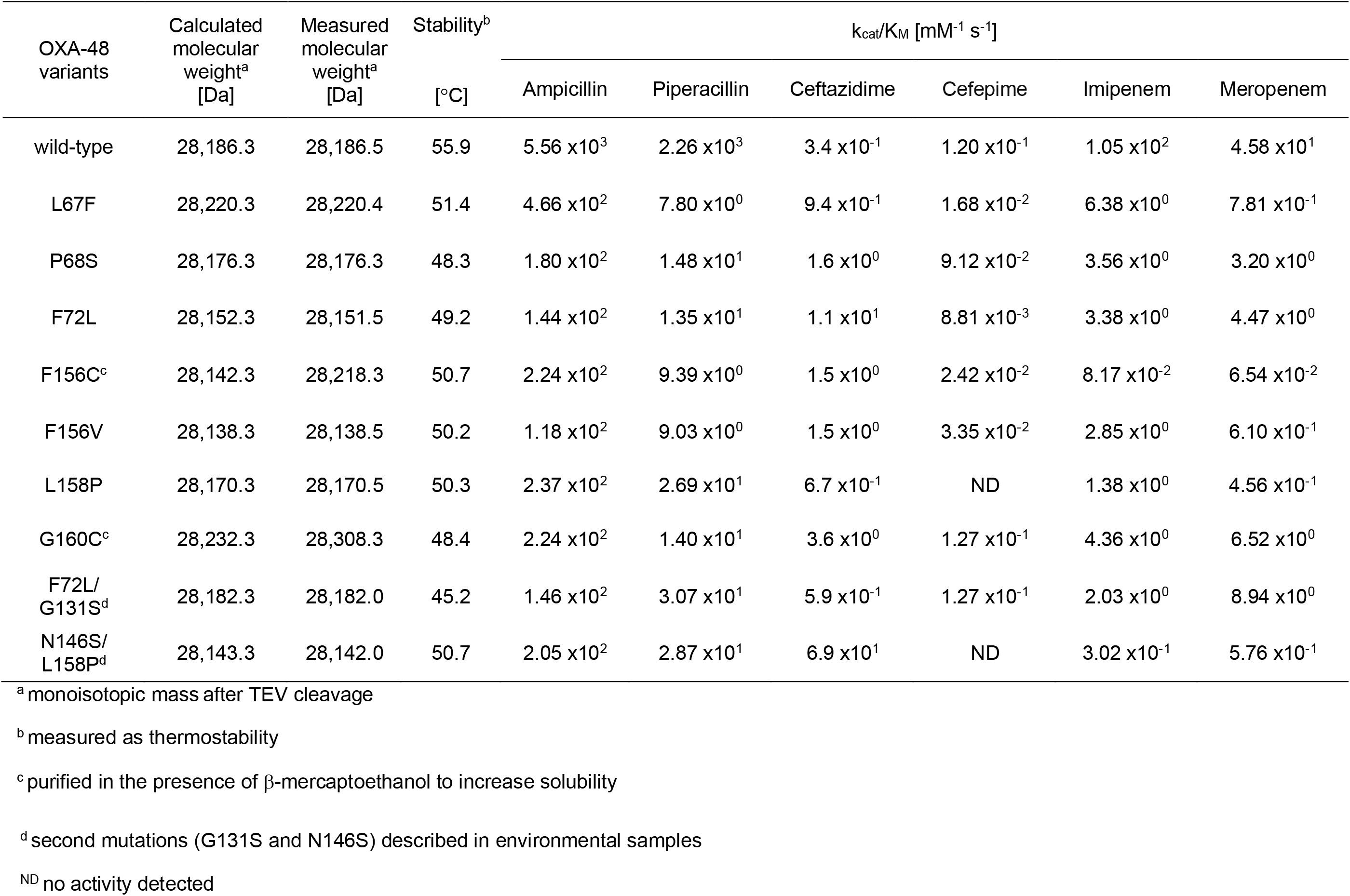
Overview of enzyme kinetic values, molecular weight and thermal stability of OXA-48 and variants.

We determined their catalytic efficiencies (k_cat_/K_M_) towards a panel of β-lactams and found that they were in-line with the antimicrobial susceptibility data. Towards ceftazidime, k_cat_/K_M_ values were increased by 2- to 31-fold (Table 2) for all variants compared to wild-type OXA-48. Moreover, all variants exhibited strongly reduced activity (up to several magnitudes) against penicillins (ampicillin and piperacillin) as well as a towards carbapenems (meropenem and imipenem). To test for cross-activity against 4^th^ generation cephalosporins, we determined the catalytic efficiencies against cefepime. Also here, we found that the OXA-48 variants tended to display k_cat_/K_M_ values several magnitudes lower than the wild-type OXA-48 (Table 2).

Functional mutations within serine β-lactamases have frequently been described to decrease the thermostability (Fröhlich et al., 2019; Mehta et al., 2015; Thomas et al., 2010). Indeed, compared to wild-type OXA-48, all single amino acid changes were deleterious with respect to thermostability, which decreased by 4.5 to 7.6°C (Table 2). F72L/G131S exhibited the lowest melting temperature with a decrease of 10.7°C. Generally, we found the following order for the thermal stability OXA-48 > L67F > F156C = N146S/L158P > L158P = F156V > F72L > G160C = P68S > F72L/G131S.

### P68S, F72L and L158P increase the loop flexibility within OXA-48

We found that single amino acid changes in OXA-48 were responsible for increased catalytic activity against ceftazidime. To understand the underlying structural changes allowing these OXA-48 variants to hydrolyse ceftazidime more efficiently, we first mapped all amino acid changes onto the structure of OXA-48 showing that they clustered around the α3-helix (L67F, P68S and F72L) and the Ω-loop (F156C, F156V, L158P and G160C) (Figure 2C). Second, MD simulations were performed on a sub-set of variants (P68S, F72L and L158P) with covalently bound ceftazidime in their active site. Our previous study showed that an amino acid change at position 68 (P68A) decreases ceftazidime susceptibility in OXA-48 (Fröhlich et al., 2019). Additionally, positions 72 and 158 were selected due to amino acid changes recently identified in environmental samples (F72I and L158P). Changes in enzyme flexibility were analysed by calculating root mean square fluctuations (RMSF) for the backbone atoms in the Ω- and β5-β6 loops and compared to wild-type OXA-48 and the ceftazidimase OXA-163 (only Ω-loop, due to the shortened β5-β6 loop).

For the Ω-loop, P68S displayed very similar RMSF values relative to OXA-48; however, F72L and L158P showed increased flexibility in this region displaying even higher RMSF values than OXA-163 (Figure S3). Notably, the L158P substitution increased fluctuations specifically for residues N152 to S155. V153 demonstrated the largest overall shift in RMSF values with an increase of 0.7 Å, when compared to OXA-48. For the β5-β6 loop, all variants exhibited an increase in fluctuations especially for the residues T213 to E216 (Figure S3).

Possible changes in intramolecular interactions due to the amino acid changes P68S, F72L and L158P were also studied from MD simulations. For P68S, an H-bond was observed between the hydroxyl groups of S68 and Y144 (Figure 2D). However, no other apparent structural changes near the active site were directly observed, and therefore the effect of P68S on the dynamical nature remains subtle.

In wild-type OXA-48, W157 in the Ω-loop stacks with both F72 and F156 (Figure 2D). Consequently, the lack of this interaction in F72L likely increases the flexibility of W157, which is reflected by a 0.2 Å increase in calculated RMSF (Figure S3). Furthermore, the wild-type Ω-loop displays an organised H-bond network, which extends to R128 and R214 on either side (Figure 2D). We found that L158P is likely affecting this network by disrupting the interactions to S155 and D159. (Figure 2D). Consequently, the salt bridge between R128 and D154 was found to be weakened as its presence was reduced from 87% to 43% of the simulation time. The loss of the backbone H-bond between L158 and S155, in the proline variant (L158P), has a knock-on effect on the rest of the loop, making it more flexible and likely to better accommodate bulkier β-lactam substrates such as ceftazidime.

Aside from flexibility and changes in amino acid interactions, possible further effects on the overall enzyme dynamics were inspected by performing principal component (PC) analysis on the combined MD trajectories (using the Cα-atom positions). The overall sampling of conformational space is highly similar for OXA-48 and the three variants (as indicated by histograms of the obtained PCs for all four enzymes, see Figure S4). There are no specific large conformational changes or coordinated loop movements induced by the mutations (with the first five PCs needed to cover ~75% of the variance in the data, Figure S5). Some differences between variants and wild-type OXA-48 were observed particularly for PCs that primarily involve movement of loops, including those surrounding the active site, further indicating that the mutations introduce small changes in loop dynamics (Figures S4 and S6).

### Release of ceftazidime from OXA-48 involves the β5-β6-loop

To investigate substrate binding, all OXA-48 variants were crystalised and soaked with ceftazidime. We were able to solve the crystal structure of L67F to 1.9 Å with four chains (A to D) in the asymmetric unit (space group P2_1_2_1_2_1_) which were arranged into two dimers (chains A/C and B/D). Chain C and D carried a hydrolysed ceftazidime molecule approximately 9 Å away from the active site S70. The R1-group of ceftazidime including the dihydrothiazine ring demonstrated clear electron density (2Fo-Fc), however, no electron density was observed for the R2-ring (Figure 3, top).

**Figure 3.**
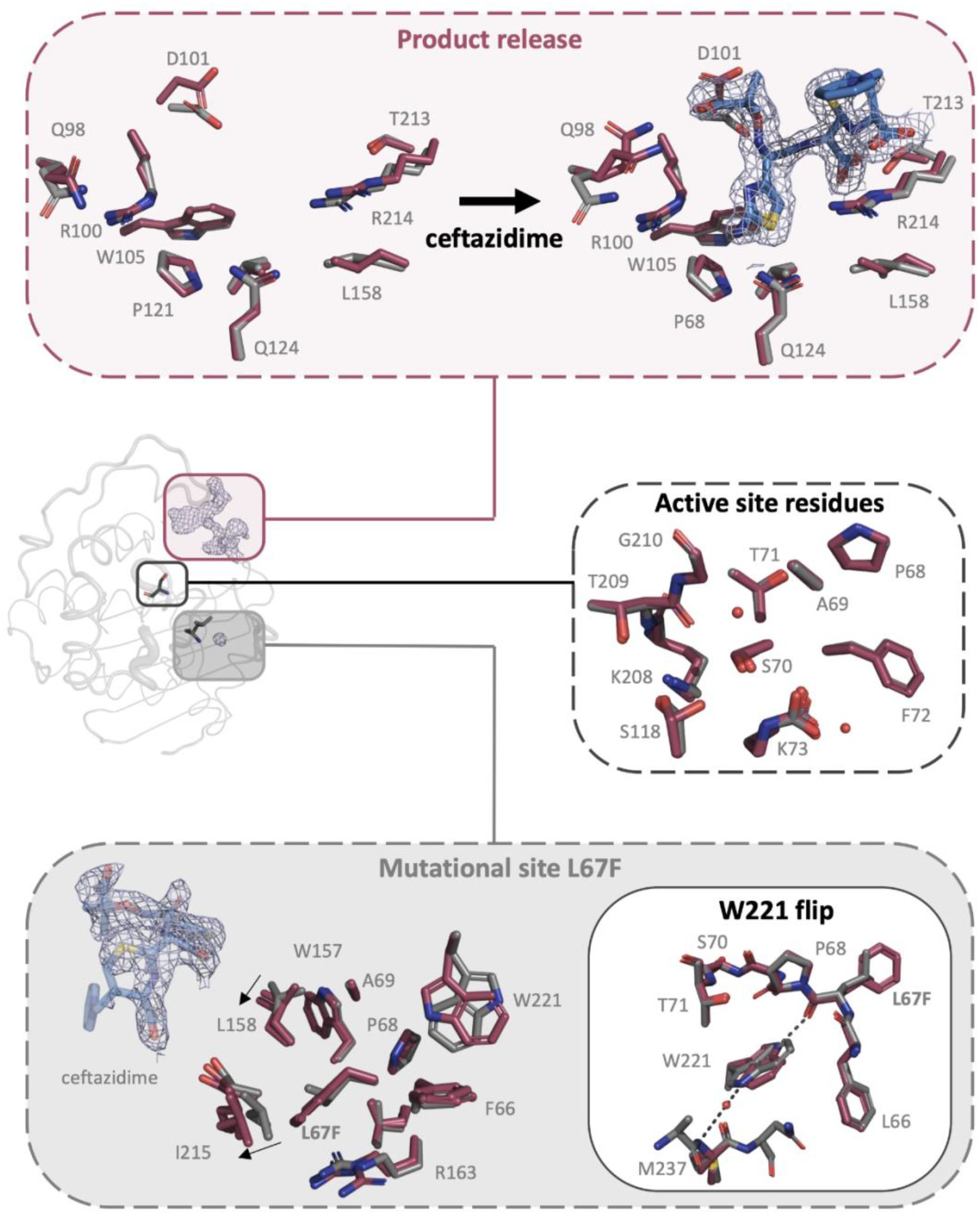
Product release (top panel), active site (middle) and the mutational site (bottom panel) of L67F (red) compared to the wild-type structure of OXA-48 shown in grey (PDB no. 3HBR) (Docquier et al., 2009). The crystal structure of L67F was solved to 1.9 Å and displayed hydrolysed ceftazidime ~ 9 Å away from the active site S70. Top panel: Binding pocket of L67F without (left, chain C) and with ceftazidime (right, chain A) compared to wild-type OXA-48. For ceftazidime, no 2Fo-Fc electron density was detected for the R2 group. Middle panel: Superimposition of the first shell residues of L67F (chain A) around the active site S70, compared the wild-type structure. Bottom panel: Investigation of the mutational site, shown as first shell residues around L67F (both chain A and C), compared to wild-type OXA-48. Displacement of L158 (1 Å) and I215 (2 Å) in the L67F structure are indicated with arrows. W221 was flipped by 180° in the L67F structure.

We first investigated the binding of ceftazidime to the L67F variant. Here, we found Q98, R100, D101, W105, V120, P121, Q124, L158, T213 and R214 to be involved in, what we believe reflects, the product release (Figure 3, top). D101, Q124, T213 and R214 were found to interact with ceftazidime *via* H-bonds. R214 was further involved in ionic interactions with the two carboxylic acid groups of ceftazidime (Figure S7).

Second, we investigated the active site architecture, including the first shell residues around S70. Chain A was therefore superimposed onto a wild-type structure of OXA-48 (PDB no. 3HBR) (Docquier et al., 2009). As expected, superimposition resulted in low root mean square deviations of 0.21 Å. We found that K73 was carboxylated and that all first shell residues nicely aligned with the wild-type structure (Figure 3, middle).

Third, we investigated the mutational site around L67F that is located “below” the active site (Figure 3, bottom). We found F66, P68, A69, W157, L158, R163, I215 and W221 to be directly interacting with amino acid position 67. In the L67F variant, both with and without ceftazidime, L158 and I215 were shifted by 1 to 2 Å, respectively. In addition, we found W221 to be flipped by 180°. While in the wild-type structure, the W221 side chain forms a water mediated H-bond to the backbone nitrogen of M237, in L67F, W221 formed a H-bond to the main chain of F67 (Figure 3).

## DISCUSSION

Here, we asked if sub-MIC of the clinically relevant β-lactam ceftazidime could affect the evolution of the contemporary, globally circulating carbapenemase OXA-48. To test this, we evolved an OXA-48 producing *E. coli* strain in the presence of one quarter of its ceftazidime MIC. The identification of seven single variants of OXA-48, conferring only marginal changes in susceptibility (Figure 2B, Table S2), demonstrates that the exposure to sub-MIC of ceftazidime drives the emergence of cryptic *bla*_OXA-48_ genetic diversity. Thus, our data provide further support for the proposed “compartment hypothesis” (Baquero & Negri, 1997; Baquero et al., 1997, 1998a, 1998b), where low-grade selection promotes cryptic genetic variation that could act as stepping-stones towards full clinical antibiotic resistance (Baier et al., 2019; Zheng et al., 2019). Notably, even though all seven single OXA-48 variants largely displayed, from a clinical microbiology perspective, neglectable changes in ceftazidime susceptibility, competition experiments revealed strong beneficial fitness effects (Figure 2B). Taken together with earlier work, using reconstructed TEM-1 variants from clinical samples (Negri et al., 2000), our data underscore the significance of divergent evolution and selection of genetic variation imposed by sub-MIC of β-lactams.

To further our understanding of how the detected single amino changes affect the structure-activity relationship of OXA-48, we first measured enzyme kinetics (Table 2). The catalytic efficiency mirrored the observed changes in susceptibility towards β - lactams at the cellular level and confirmed our previous findings that mutational changes increasing ceftazidime activity comes with a functional trade-off against penicillins and carbapenems (Fröhlich et al., 2019). Structurally all amino acid changes clustered either around the active site S70 (L67F, P68S, F72L) or within the Ω-loop (F156C, F156V, L158P, G160C; Figure 2C).

In wild-type OXA-48, the Ω-loop interacts with the β5-β6 loop *via* a salt bridge mediated by D159-R214 maintaining a closed conformation of the active site (Docquier et al., 2009). MD simulations performed on a sub-set of variants revealed that F72L and L158P weaken the interaction between these loops resulting in increased structural flexibility (Figure S3). We postulate that these changes aid the hydrolysis of bulkier substrates like ceftazidime but result in decreased activity towards penicillins and carbapenems. Indeed, mutations affecting the salt bridge are associated with reduced carbapenemase activity presumably due to increased loop flexibility (Oueslati et al., 2020).

In clinical OXA-48-like variants (e.g. OXA-163, OXA-247 and OXA-405) with increased ceftazidime activity, larger structural variations (deletions in combination with single point mutations) within and around the β5-β6 loop have been reported (Dortet et al., 2015; Gomez et al., 2013; Poirel et al., 2011). However, also single amino acid changes structurally close to the Ω- and β5-β6 loops have been shown to slightly elevate the catalytic efficiency towards ceftazidime (E125Y in OXA-245 and V120L in OXA-519) (Dabos et al., 2018; Lund et al., 2017). The significance of the Ω- and β5-β6 loops for the substrate profile is not limited to OXA-48 (Dabos et al., 2020; De Luca et al., 2011). These loops have been shown to impact substrate profiles for other clinically relevant β-lactamases including TEM and KPC (Levitt et al., 2012; Stojanoski et al., 2015; Venditti et al., 2019). In addition, comparable modes of action have been hypothesised for the OXA-10-like variants OXA-145 and OXA-147 exhibiting L158Δ and W157L, respectively (according to OXA-48 numbering) (Baurin et al., 2009; Fournier et al., 2010; Meziane-Cherif et al., 2016).

We were able to solve the structure of the OXA-48 variant L67F revealing a binding site for ceftazidime, approximately 9 Å away from the active site residue S70, involving interaction with the above described β5-β6 loop (Figure 3) and R214 in particular. Since ceftazidime was hydrolysed, we hypothesise that this may reflect the product release process (Figure S7).

Taken together, combining experimental evolution and structure-activity relationships allowed us to identify and characterize single step mutations with cryptic resistance that yet demonstrated significant fitness effects and structural changes. Our data show that to understand the evolutionary potential of standing genetic diversity, susceptibilities characterised solely by traditional MIC measurements provide too low resolution.

We acknowledge that our study is not without limitations as that, despite strong fitness effects, none of the variants went to fixation in any of the populations (Figure 1). We argue that there can be at least two reason for this. First, we have focused solely on OXA-48 mutations and it is clear from our data that the evolution at sub-MIC also selected for other potential resistance mechanisms. These mechanisms could lead to potential wide-spread epistatic interactions that would slow down any fixation process (de Visser & Rozen, 2006; Gullberg et al., 2011; Shields, Chen, et al., 2017; Shields, Nguyen, et al., 2017; Westhoff et al., 2017). Second, it has been shown that β-lactamase producers can detoxify their environment allowing co-existence of genotypes with different susceptibilities resulting in clone-frequency equilibria (Yurtsev et al., 2013).

Our work sheds light on the evolution of β-lactamases and their selection dynamics towards altered substrate profiles. This is supported by recent studies reporting environmental contamination of cephalosporins at concentrations similar to those applied here (Ribeiro et al., 2018; Watkinson et al., 2009). Moreover, OXA-48 variants, with the same or similar amino acid changes as identified and characterized here, have been reported in environmental samples (Naas et al., 2017; Tacao et al., 2015). We speculate that the identified mutations are only first step mutations towards full clinical ceftazidime resistance mediated by OXA-48. However, more studies are needed to fully understand the complete fitness landscape of OXA-48 and other carbapenemases.

## METHODS

### Media, chemicals and strains

Mueller Hinton (MH) agar and broth were purchased from Thermo Fisher Scientific (East Grinstead, UK). Luria-Bertani (LB) broth, LB agar, yeast extract, agar, terrific broth, ampicillin, amoxicillin, cefepime, ceftazidime, chloramphenicol, imipenem, meropenem, piperacillin, 2,3,5 tri-phenyl tetrazolium, sodium chloride and maltose were obtained from Sigma-Aldrich (St. Louis, MO, USA). Tryptone was obtained from Oxoid (Hampshire, UK) All strains used and constructed within this study are listed in Table S1.

### Sub-MIC evolution

MP13-06, previously constructed and tested (Fröhlich et al., 2019), was evolved by serial passaging without selection pressure and at 0.25xMIC (0.06 mg/L) of ceftazidime for 300 generations. In short, bacterial suspensions were grown at 37°C, 700 rpm on a plate shaker (Edmund Bühler, Bodelshausen, Germany) in 1 mL MH broth to full density and passaged every 12 h with a bottleneck of 1:100. The evolution was performed in triplicates.

### Dose-response curves and susceptibility testing

Dose-response curves were obtained initially and after every 50 generations for the whole evolved populations. Cultures were grown to full density and diluted in 0.9% saline to 10^6^ CFU/mL. 384 well plates (VWR, Radnor, PA, USA) were inoculated with 10^5^ CFU and increasing concentrations of ceftazidime ranging from 0 to 32 mg/L. Plates were statically incubated for 20 h at 37°C. The optical density at 600 nm (OD_600_) was measured with a microtiter plate reader (Biotek Instruments, Winooski, VT, USA) and dose-response curves including IC_50_ values were calculated using GraphPad Prism 9.0 (GraphPad Software, San Diego, CA, USA). In this set-up, the ceftazidime MIC was determined by measuring the OD_600_ as the first well with an optical density, comparable to the negative control.

For MIC measurements against other β-lactams than ceftazidime, in-house designed and premade Sensititre microtiter plates (TREK Diagnostic Systems/Thermo Fisher Scientific, East Grinstead, UK) were loaded with 10^5^ CFU. The plates were incubated statically for 20 h at 37°C. All susceptibility tests were performed in at least two biological replicates.

### Determination of clones with altered ceftazidime susceptibility

To determine clones exhibiting decreased ceftazidime susceptibility, we plated 10^7^ cells from every 50^th^ generation on MH agar without and with 1 mg/L ceftazidime. The plates were incubated for 24 h at 37°C. Clone frequencies were determined as the ratio between colonies found on selective *versus* non-selective plates. About 50 pre-selected colonies were subjected to susceptibility testing, as described above. For creating the boxplots, a probability function was calculated based on MIC per replicate. Since the MIC were determined in 2-fold steps, we generated smoother boxplots by creating 1000 random measurements per generation. We have done so by drawing a random number between the log2 (MIC values) and log2 (MIC values) +1 a 1000 x f_mic, where f_mic is the fraction of the population. All calculations were done in Mathematica 11.0 (Wolfram Research, Champaign, IL, USA).

### Strain construction

For functional resistance profiles, wild-type *bla*_OXA-48_ and allele variants were sub-cloned into the high copy number vector pCR-blunt II-TOPO vector (Invitrogen, Carlsbad, CA, USA) and expressed in *E. coli* TOP10 (Invitrogen). For wild-type TOPO-*bla*_OXA-48_, the construction has been described previously (Fröhlich et al., 2019). Point mutations were inserted by using the Quick-change II kit for site directed mutagenesis (Agilent Biosciences, Santa Clara, CA, USA), TOPO-*bla*_OXA-48_ as a template and the respective primers (Table S3). The double mutants TOPO-*bla*_OXA-48_-F72L/G131S and TOPO-*bla*OXA-48-N146S/L158P were created by inverse PCR using Phusion polymerase (New England Biolabs, Ipswich, MA, USA) and TOPO-*bla*_OXA-48_-F72L or TOPO-*bla*_OXA-48_-L158P as a template, respectively. PCR products were 5′-phosphorylated with polynucleotide kinase (Thermo Fisher Scientific, Waltham, MA, USA), and circularised using T4 DNA ligase (Thermo Fisher Scientific). Transformants were selected on LB agar plates containing 50 or 100 mg/L ampicillin. *Bla*_OXA-48_ was Sanger sequenced (BigDye 3.1 technology, Applied Biosystems, Foster City, CA, USA) using M13 primers (Thermo Fisher Scientific) (Table S3).

For expression in a low copy number vector (pUN), we PCR-amplified a segment containing the p15A origin of replication and the *cat* chloramphenicol resistance gene of the pACYC184 vector using the primers cat-r and p15A46 (Table S3). To obtain the *bla*_OXA-48_ inserts, the pCR-blunt II-TOPO constructs (see above) were used as templates. We amplified the *bla*_OXA-48_ genes by using the primers OXA-48-pro-f, containing the constitutive artificial CP6 promoter (Jensen & Hammer, 1998), and preOXA-48B (Table S3) (Samuelsen et al., 2013). These PCR products were 5′-phosphorylated with polynucleotide kinase (Thermo Fisher Scientific) and then blunt ligated with the amplified vector backbone. The resulting in pUN-*bla*_OXA-48_ vector and the corresponding variants were transformed into *E. coli* DH5α and plated on LB agar containing chloramphenicol (25 mg/L). Genotypes of selected clones were confirmed by Sanger sequencing (BigDye 3.1 technology, Applied Biosystems) using preOXA-48A/B primers (Table S3) (Samuelsen et al., 2013).

To measure bacterial fitness, *E. coli* MG1655 Δ*malF* (MP14-23) was constructed as a competitor strain by transducing the kanamycin resistance marker from the Keio strain JW3993 with P1-vir into *E. coli* MG1655 as published (Baba & Mori, 2008; Thomason et al., 2007). The marker was then removed with the helper vector pCP20 (Datsenko & Wanner, 2000). The competitor strain MP14-23 was then transformed with pUN-*bla*_OXA-48_ and the corresponding variants (Table S1). Transformants were selected on LB plates containing 25 mg/L chloramphenicol.

For protein expression and purification, *bla*_OXA-48_ in the pDEST17 expression vector (Thermo Fisher Scientific) was mutagenized using QuickChange II site-directed mutagenesis kit as described above. *E. coli* DH5α was transformed with the DNA constructs and clones were selected on LB agar containing 100mg/L ampicillin. The vectors were isolated using a plasmid maxi kit (Qiagen, Hilden, Germany) and transformed into *E. coli* BL21 AI (Thermo Fisher Scientific). Point mutations were verified by Sanger sequencing using T7 primers (Thermo Fisher Scientific) (Table S3).

### Bacterial fitness: head-to-head competition

Strains were grown overnight in LB supplemented with chloramphenicol (25 mg/L) at 37°C and 700 rpm on a plate shaker (Edmund Bühler). For each competition, we co-inoculated ~ 1×^7^ CFU/mL of each competitor in 1 mL LB broth, supplemented with either chloramphenicol (25 mg/L) and ceftazidime (0.06 mg/L), or with chloramphenicol only. 96-deep-well plates (VWR) were incubated at 37°C and 700 rpm for 8 h. Each competition was performed in three biological replicates. Initial and final CFU/mL for both competitors were determined by differential plating on tetrazolium maltose agar (10 g/L tryptone, 5 g/L sodium chloride, 1 g/L yeast extract, 15 g/L agar, 10 g/L maltose, supplemented with 1 mL 5% 2,3,5 tri-phenyl tetrazolium chloride). Relative fitness (w) was determined according to equation 1, where *mal^+^* and *mal^-^* are respectively the *mal*^+^ and Δ*malF* strain backgrounds carrying the different pUN vectors (Table S1).

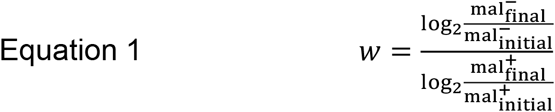

The *mal*^+^ pUN-*bla*_OXA-48_ strain (MP08-61) was competed against Δ*malF* strains carrying each of the pUN vectors encoding wild-type *bla*_OXA-48_ or single variants (Table S1). Additionally, the *mal*^+^ pUN-*bla*_OXA-48_-F72L (MP08-67) and *mal*^+^ pUN-*bla*_OXA-48_-L158P (MP08-63) strains were used as competitors respectively against strains Δ*malF* pUN-*bla*_OXA-48_-F72L/G131S (MP14-32) and Δ*malF* pUN-*bla*_OXA-48_-N146S/L158P (MP14-33), respectively. Data analysis and graphical illustrations were performed in R version 4.0.2 (RCoreTeam, 2018).

### Recombinant enzyme expression and purification

Overexpression of OXA-48 and the corresponding variants was done in terrific broth supplemented with 100 mg/L ampicillin. *E. coli* BL21 AI carrying pDEST-17-*bla*_OXA-48_ and OXA-48 variants (Table S1) were grown at 37°C and 220 rpm to an optical density of 0.4 to 0.5. Protein expression was induced with 0.1% L-arabinose (Sigma-Aldrich). Expression took place for 16 h at 15°C and 220 rpm. Harvested cells were sonicated, and recombinant proteins were purified as described previously (Fröhlich et al., 2019; Lund et al., 2016). F156C and G160C were found to be insoluble. To increase their solubility, 5 mM β-mercaptoethanol was used during the sonication process.

### Molecular mass verification

ESI-MS was performed on the purified enzymes as described previously (Fröhlich et al., 2020). In short, a buffer exchange to 0.1% formic acid (Merck Millipore, Burlington, MA, USA) was performed using centrifugal molecular cut-off filters (Merck Millipore; 10,000 Da). The protein masses were determined using an Orbitrap Fusion Lumos (Thermo Fisher Scientific). Injection was performed using an EASY-nano LC (Thermo Fisher Scientific) with a 15 cm C18 EASY-spray column. Mass calculations were done using the BioPharma Finder 3.0 protein deconvolution software (Thermo Fisher Scientific, MA, USA).

### Steady-state enzyme kinetics

Catalytic efficiencies (k_cat_/K_M_) for the recombinantly expressed enzymes were determined under steady-state conditions for ampicillin (Δξ= - 820 M^-1^ cm^-1^, 232 nm), piperacillin (Δξ= - 820 M^-1^ cm^-1^, 235 nm), ceftazidime (Δξ= - 9,000 M^-1^ cm^-1^, 260 nm), cefepime (Δξ= - 10,000 M^-1^ cm^-1^, 260 nm), imipenem (Δξ= - 9,000 M^-1^ cm^-1^, 300 nm), and meropenem (Δξ= - 6,500 M^-1^ cm^-1^, 300 nm) by measuring the initial enzymatic reaction rate. Enzyme concentrations are summarised in Table S4. All determinations were performed at least in duplicates at a final assay volume of 100 μL. UV-transparent 96 well plates (Corning, Kennebunk, ME, USA) were used. All test results were obtained at 25°C and in 0.1 M phosphate buffer (pH 7.0) supplemented with 50 mM NaHCO_3_ (Sigma Aldrich). Calculations were performed by using GraphPad Prism 9.0 (GraphPad Software).

### Thermostability

We determined the fluorescence-based protein thermostability for OXA-48, as described previously (Fröhlich et al., 2019). In short, the proteins were diluted in 50 mM HEPES (VWR), pH 7.5 supplemented with 50 mM potassium sulphate (Honeywell, NC, USA) to a final concentration of 0.2 mg/mL protein and 5x SYPRO orange (Sigma-Aldrich). A temperature gradient of 25 to 70°C (heating rate 1°C per min) was applied using a MJ minicycler (Bio-Rad, Hercules, CA, USA). All experiments were performed triplicates.

### Molecular dynamics simulations

System set-up was done as described previously (further details in the SI) (Hirvonen et al., 2020). All systems were initially briefly minimized (1000 steps of steepest descent followed by 1000 steps of conjugate gradient), heated from 50 K to 300 K in 20 ps, and then simulated for 120 ns in the NPT ensemble (saving a frame every 20 ps). Langevin dynamics were used with a collision frequency of 0.2 and a 2 fs timestep, all bonds involving hydrogens were constrained using the SHAKE algorithm. Periodic boundary conditions in explicit solvent were applied in all simulations. Five independent simulations were run per enzyme variant (for a total of 600 ns per variant), and all calculations were done with the Amber18 program package (pmemd.cuda) (Rubenstein et al., 2018) using the ff14SB force field (Maier et al., 2015) for the protein and TIP3P for water (Grand et al., 2013; Salomon-Ferrer et al., 2013). All analyses were done using cpptraj from AmberTools (Roe & Cheatham, 2013). Further computational details can be found in the supplementary material.

### Crystallization and structure determination

Crystals were grown in a 1 μL hanging drop containing 5 mg/mL enzyme and mixed 1:1 with reservoir solution containing 0.1 M Tris, pH 9.0 (Sigma-Aldrich) and 28-30% PEG mono ethylene ether 500 (Sigma-Aldrich) at 4°C. Crystals were harvested, cryoprotected by adding 15% ethylene glycol (Sigma-Aldrich) to the reservoir solution and then frozen in liquid nitrogen.

Diffraction data were collected on BL14.1 BESSY II, Berlin, Germany, at 100 K, wavelength 0.9184 Å, and the diffraction images were indexed and integrated using XDS (Kabsch, 2010). AIMLESS was used for scaling (Evans & Murshudov, 2013). When scaling the final dataset (Table S5), we aimed for high overall completeness, and CC_1/2_ > 0.5 and a mean intensity <I> above 1.0 in the outer resolution shell. The structure was solved by molecular replacement with chain A of PDB no. 5QB4 (Akhter et al., 2018) and the program Phenix 1.12 (Adams et al., 2010). Parts of the model was rebuilt using Coot (Emsley et al., 2010). Figures were prepared using PyMOL version 1.8 (Schrödinger, New York City, NY, USA). Ligand and protein interactions were calculated using Protein Contact Altas (Kayikci et al., 2018).

## AUTHOR CONTRIBUTIONS

CF, PJJ, ØS and HKSL worked out the conceptional framework. CF conducted the evolution. CF and JAG performed microbiological testing. CF, JAG and KH constructed strains. CF purified enzymes and measured kinetics. CF and HKSL solved the crystal structure. BAL performed initial molecular dynamic simulations as a proof of principle. VHAH and MWvdK designed, performed and analysed the molecular dynamic simulations. CF wrote the manuscript with contributions of all co-authors.

## Data availability/accession numbers

Atom coordinates and structure factors for the OXA-48 variant L67F are deposited in the protein data bank (PDB no. 7ASS). Data will be made available as source files.

## ACKNOWDLEGMENTS

We are very grateful to Linus Sandegren, Uppsala University, Sweden, and Francisco Dionísio, University of Lisbon, Portugal, for providing strains. We thank Karina Xavier (The Instituto Gulbenkian Ciência, Portugal) for providing the P1 phage. Provision of beam time at BL14.1 at Bessy II, Berlin, Germany is highly valued. Hanna-Kirsti S. Leiros was supported by Norwegian Research Council (273332/2018). Pål Jarle Johnsen was supported by Northern Norway Regional Health Authority, UiT The Arctic University of Norway (Project SFP1292-16), and JPI-EC-AMR (Project 271176/H10). Viivi H.A. Hirvonen was supported by the UK Medical Research Council (MR/N0137941/1). Marc W. van der Kamp is a BBSRC David Phillips Fellow and thanks the Biotechnology and Biological Sciences Research Council for funding (BB/M026280/1). Simulations were performed using the computational facilities of the Advanced Computing Research Centre, University of Bristol. We would also like to extend our thanks to François Pierre Alexandre Cléon, Vidar Sørum, Antal Martinecz and Alexander Wessel for their help.

## COMPETING INTERESTS

None to declare.

## SUPPLEMENTARY MATERIAL

**Supplementary information:** molecular dynamics simulations

### Supplementary tables

**Table S1.** Strains used and constructed in this study

**Table S2.** Ceftazidime susceptibility (MIC and IC50) measurements (mg/L) of OXA-48 and variants expressed from the low copy number vector pUN in E. coli MG1655ΔmalF (MP08-23). Susceptibility was determined based on a minimum of 2 biological replicates. The 95% confidence interval [CI95%] were calculated for the IC50 values.

**Table S3.** Primers used in the study

**Table S4.** Enzyme concentrations (nM) for steady-state kinetics

**Table S5.** X-ray data collection and refinement statistics for the OXA-48 variant L67F in complex with hydrolysed ceftazidime. Values in parenthesis are for the highest resolution shell.

### Supplementary figures

**Figure S1.** Head-to-head competitions, between *E. coli* MG1655 *mal*^+^ and MG1655 Δ*malF* expressing wild-type and allele variants of OXA-48, conducted without (grey) and at sub-MIC (red) of ceftazidime. While expression without selection pressure was neutral for all alleles, at sub-MIC, all allele variants showed fitness benefits over the wild-type allele. The dots represent biological replicates and significantly different averages, compared to OXA-48 in the presence of ceftazidime (0.06 mg/L), are marked with * (P< 0.05), ** (P<0.01) and *** (P<0.001).

**Figure S2.** Head-to-head competitions between MG1655 expressing F72L versus F72L/G131S and L158P versus N146S/L158P. G131S and N146S did not improve bacterial fitness at sub-MIC ceftazidime. The dots represent biological replicates. Significant differences are indicated as * representing P< 0.05.

**Figure S3.** Root mean square fluctuations (RMSFs) for the clinical variants OXA-48 and OXA-163 as well as for a sub-set of OXA-48 variants: P68S, F72L and L158P. RMSFs for residues in the Ω-loop (A) and for the β5-β6 loop (B).

**Figure S4**. Normalized histograms of PC1-PC5 (A to E) for OXA-48 and the variants P68S, F72L and L158P. The histograms are calculated using 200 bins per enzyme.

**Figure S5.** Cumulative variance covered by the ten principal components.

**Figure S6.** Dynamic differences between wild-type OXA-48, P68S, F72L and L158P captured by PC1 (A) and PC5 (B). Arrows indicating Cα-movement in PC1 (A) and PC5 (B), arrow direction and size indicating direction of the eigenvector and magnitude of the corresponding eigenvalue (arrows only shown for atoms with eigen values > 2.5 Å).

**Figure S7:** Schematic representation of hydrolysed ceftazidime in front of the active site of the OXA-48 variant L67F (based on PDB no. 7ASS). The ceftazidime side chains R1 and R2 are labelled and marked. For R2, no electron density and therefore no interactions were detected. Hydrogen bonds from ceftazidime to D101, Q124, T213 and R214 are represented with dashed lines. The salt bridges to R214 are indicated with arrows.

## Notes

### Competing Interest Statement

The authors have declared no competing interest.

