## Supplementary material for "Cryptic β-lactamase evolution is driven by low β-lactam concentrations"

**Supplementary information:** molecular dynamics simulations

System set-up

Acylenzyme of OXA-48 with covalently bound ceftazidime was originally built using the structure of OXA-48 with imipenem (PDB no. 5QB4) (Akhter et al., 2018), and replacing imipenem as found in the OXA-48 co-complex with ceftazidime (PDB no. 6Q5F) (Fröhlich et al., 2019), which ensured keeping the Ω-loop ordered (as found in the apoenzyme). For OXA-163, the same ceftazidime binding pose was combined with apoenzyme crystal structure (PDB no. 4S2L) (Stojanoski et al., 2015). All OXA-48 variants were modelled using the mutagenesis tool in PyMOL, choosing the rotamer with least steric clashes with surrounding atoms. Enzymes were solvated in a 10 Å cubic box of TIP3P water, any existing net charges were neutralized by replacing bulk water molecules with counter ions. Non-standard residues (carboxylated K73, ceftazidime acylenzyme) were parametrized using the RED Server, have been published previously (Hirvonen et al., 2020).

RMSF calculations

Root mean square fluctuation (RMSF) calculations were performed by first calculating an average structure from the MD simulation, aligning the trajectory against the average structure excluding four residues from both the N- and C- terminal ends as well as the loop residues of interest (G145-I164 for the Ω-loop and T213-I219 for the β5-β6-loop), and then calculating RMSF for the corresponding loop mainchain heavy atoms (CA, C, N, O). The first 20 ns were excluded from all analyses to allow time for system equilibration.

Principal Component (PC) Analysis

PC analysis was performed on Cα-atoms (excluding four residues from both the N- and C-terminal ends), using all trajectories of OXA-48 and variants together (5 independent simulations of 120 ns per variant). A total of ten principal components (PCs) were calculated. PCs were projected onto trajectory data to compare the sampled PC values between enzymes, and normalized frequency histograms of values for PC1-PC5 are presented in Figure S4.

### Supplementary tables

**Table S1.** Strains used and constructed in this study

| *E. coli* Strain(s) | Comment | Reference |
|---|---|---|
| MP13-01/ MP08-01 | K12 MG1655 | Uppsala/Lisbon University |
| MP13-06[a] | MP13-01 transformed with p50579417_3_OXA-48 | (Fröhlich et al., 2019) |
| MP13-34 | MP13-06 evolved in MH broth, population 1: 50 generations | this study |
| MP13-35 | MP13-06 evolved in MH broth, population 1: 100 generations | this study |
| MP13-36 | MP13-06 evolved in MH broth, population 1: 150 generations | this study |
| MP13-37 | MP13-06 evolved in MH broth, population 1: 200 generations | this study |
| MP13-38 | MP13-06 evolved in MH broth, population 1: 250 generations | this study |
| MP13-39 | MP13-06 evolved in MH broth, population 1: 300 generations | this study |
| MP13-40 | MP13-06 evolved in MH broth, population 2: 50 generations | this study |
| MP13-41 | MP13-06 evolved in MH broth, population 2:100 generations | this study |
| MP13-42 | MP13-06 evolved in MH broth, population 2:150 generations | this study |
| MP13-43 | MP13-06 evolved in MH broth, population 2: 200 generations | this study |
| MP13-44 | MP13-06 evolved in MH broth, population 2: 250 generations | this study |
| MP13-45 | MP13-06 evolved in MH broth, population 2: 300 generations | this study |
| MP13-46 | MP13-06 evolved in MH broth, population 3: 50 generations | this study |
| MP13-47 | MP13-06 evolved in MH broth, population 3: 100 generations | this study |
| MP13-48 | MP13-06 evolved in MH broth, population 3: 150 generations | this study |
| MP13-49 | MP13-06 evolved in MH broth, population 3: 200 generations | this study |
| MP13-50 | MP13-06 evolved in MH broth, population 3: 250 generations | this study |
| MP13-51 | MP13-06 evolved in MH broth, population 3: 300 generations | this study |
| MP13-52 | MP13-06 evolved in ceftazidime, population 4: 50 generations | this study |
| MP13-53 | MP13-06 evolved in ceftazidime, population 4: 100 generations | this study |
| MP13-54 | MP13-06 evolved in ceftazidime, population 4: 150 generations | this study |
| MP13-55 | MP13-06 evolved in ceftazidime, population 4: 200 generations | this study |
| MP13-56 | MP13-06 evolved in ceftazidime, population 4: 250 generations | this study |
| MP13-57 | MP13-06 evolved in ceftazidime, population 4: 300 generations | this study |
| MP13-58 | MP13-06 evolved in ceftazidime, population 5: 50 generations | this study |
| MP13-59 | MP13-06 evolved in ceftazidime, population 5: 100 generations | this study |

| | | |
|---|---|---|
| MP13-60 | MP13-06 evolved in ceftazidime, population 5: 150 generations | this study |
| MP13-61 | MP13-06 evolved in ceftazidime, population 5: 200 generations | this study |
| MP13-62 | MP13-06 evolved in ceftazidime, population 5: 250 generations | this study |
| MP13-63 | MP13-06 evolved in ceftazidime, population 5: 300 generations | this study |
| MP13-64 | MP13-06 evolved in ceftazidime, population 6: 50 generations | this study |
| MP13-65 | MP13-06 evolved in ceftazidime, population 6: 100 generations | this study |
| MP13-66 | MP13-06 evolved in ceftazidime, population 6: 150 generations | this study |
| MP13-67 | MP13-06 evolved in ceftazidime, population 6: 200 generations | this study |
| MP13-68 | MP13-06 evolved in ceftazidime, population 6: 250 generations | this study |
| MP13-69 | MP13-06 evolved in ceftazidime, population 6: 300 generations | this study |
| | For MIC measurements (high copy number vector): | |
| MP13-04 | recipient strain for pCR-blunt II-TOPO | Invitrogen |
| MP13-11 | MP13-04, transformed with pCR-blunt II-*bla*$_{OXA-48}$ | (Fröhlich et al., 2019) |
| MP13-21 | MP13-04, transformed with pCR-blunt II-*bla*$_{OXA-48}$-L67F | this study |
| MP13-16 | MP13-04, transformed with pCR-blunt II-*bla*$_{OXA-48}$-P68S | this study |
| MP13-14 | MP13-04, transformed with pCR-blunt II-*bla*$_{OXA-48}$-F72L | this study |
| MP13-17 | MP13-04, transformed with pCR-blunt II-*bla*$_{OXA-48}$-F156C | this study |
| MP13-18 | MP13-04, transformed with pCR-blunt II-*bla*$_{OXA-48}$-F156V | this study |
| MP13-15 | MP13-04, transformed with pCR-blunt II-*bla*$_{OXA-48}$-L158P | this study |
| MP13-19 | MP13-04, transformed with pCR-blunt II-*bla*$_{OXA-48}$-G160C | this study |
| MP13-33 | MP13-04, transformed with pCR-blunt II-*bla*$_{OXA-48}$-F72L/G131S | this study |
| MP13-20 | MP13-04, transformed with pCR-blunt II-*bla*$_{OXA-48}$-N146S/L158P | this study |
| | For dose-response measurements and head-to-head competitions (low copy number vector): | |
| JW3393 | K12 BW25113, *ΔmalF*729::*kan* | (Baba et al., 2006) |
| MP14-23 | MG1655 *ΔmalF* constructed based on MP08-01 (*mal*$^{+}$) | this study |
| MP08-61 | MP08-01 with pUN-*bla*$_{OXA-48}$ | this study |
| MP14-24 | MP14-23 with pUN-*bla*$_{OXA-48}$ | this study |
| MP14-29 | MP14-23 with pUN-*bla*$_{OXA-48}$-L67F | this study |
| MP14-26 | MP14-23 with pUN-*bla*$_{OXA-48}$-P68S | this study |
| MP14-27 | MP14-23 with pUN-*bla*$_{OXA-48}$-F72L | this study |
| MP14-30 | MP14-23 with pUN-*bla*$_{OXA-48}$-F156C | this study |
| MP14-31 | MP14-23 with pUN-*bla*$_{OXA-48}$-F156V | this study |

| | | |
|---|---|---|
| MP14-25 | MP14-23 with pUN-*bla*$_{OXA-48}$-L158P | this study |
| MP14-28 | MP14-23 with pUN-*bla*$_{OXA-48}$-G160C | this study |
| MP14-32 | MP14-23 with pUN-*bla*$_{OXA-48}$-F72L/G131S | this study |
| MP08-67 | MP08-01 with pUN-*bla*$_{OXA-48}$-F72L | this study |
| MP14-33 | MP14-23 with pUN-*bla*$_{OXA-48}$-N146S/L158P | this study |
| MP08-63 | MP08-01 with pUN-*bla*$_{OXA-48}$-L158P | this study |
| | For protein expression and purification: | |
| MP13-02 | BL21 AI recipient for pDEST17 expression vector | Invitrogen |
| MP13-23 | MP13-02, transformed with pDEST17-*bla*$_{OXA-48}$ | (Fröhlich et al., 2019) |
| MP13-24 | MP13-02, transformed with pDEST17-*bla*$_{OXA-48}$-L67F | this study |
| MP13-25 | MP13-02, transformed with pDEST17-*bla*$_{OXA-48}$-P68S | this study |
| MP13-26 | MP13-02, transformed with pDEST17-*bla*$_{OXA-48}$-F72L | this study |
| MP13-27 | MP13-02, transformed with pDEST17-*bla*$_{OXA-48}$-F156C | this study |
| MP13-28 | MP13-02, transformed with pDEST17-*bla*$_{OXA-48}$-F156V | this study |
| MP13-29 | MP13-02, transformed with pDEST17-*bla*$_{OXA-48}$-L158P | this study |
| MP13-30 | MP13-02, transformed with pDEST17-*bla*$_{OXA-48}$-G160C | this study |
| MP13-31 | MP13-02, transformed with pDEST17-*bla*$_{OXA-48}$-F72L/G131S | this study |
| MP13-32 | MP13-02, transformed with pDEST17-*bla*$_{OXA-48}$-N146S/L158P | this study |

[a] previously named as MP101(Fröhlich et al., 2019)

**Table S2.** Ceftazidime susceptibility (MIC and $IC_{50}$) measurements (mg/L) of OXA-48 and variants expressed from the low copy number vector pUN in *E. coli* MG1655*ΔmalF* (MP08-23). Susceptibility was determined based on a minimum of 2 biological replicates. The 95% confidence interval [CI95%] were calculated for the $IC_{50}$ values.

| | **MP08-01** | **MP08-61 wild-type OXA-48** | **MP14-29 L67F** | **MP14-26 P68S** | **MP14-27 F72L** | **MP14-30 F156C** | **MP14-31 F156V** | **MP14-25 L158P** | **MP14-28 G160C** | **MP14-32 F72L/G131S** | **MP14-33 N146S/L158P** |
|---|---|---|---|---|---|---|---|---|---|---|---|
| MIC | 0.5 | 0.5 | 0.5 | 1 | 1 | 1 | 2 | 1 | 0.5 | 0.5 | 1 |
| $IC_{50}$ [CI95%] | 0.045 [0.033 to 0.061] | 0.053 [0.039 to 0.070] | 0.074 [0.057 to 0.096] | 0.160 [0.128 to 0.201] | 0.167 [0.136 to 0.205] | 0.161 [0.135 to 0.192] | 0.191 [0.153 to 0.239] | 0.176 [0.142 to 0.219] | 0.114 [0.087 to 0.148] | 0.078 [0.062 to 0.097] | 0.220 [0.164 to 0.295] |

**Table S3.** Primers used in the study

| Name[a] | | 5´-sequence-3´ | Reference |
|---|---|---|---|
| preOXA-48 | FP (A) | TATATTGCATTAAGCAAGGG | (Samuelsen et al., 2013) |
| | RP (B) | CACACAAATACGCGCTAACC | |
| M13 | FP | GTAAAACGACGGCCAG | Thermo Fisher Scientific |
| | RP | CAGGAAACAGCTATGAC | |
| T7 | FP | TAATACGACTCACTATAGGG | Thermo Fisher Scientific |
| | RP | GCTAGTTATTGCTCAGCGG | |
| F67L | FP | GGGCGAACCAAGCATTTTTTCCCGCATCTACC | this study |
| | RP | GGTAGATGCGGGAAAAAATGCTTGGTTCGCCC | |
| P68S | FP | CGGGCGAACCAAGCATTTTTATCCGCATCTACC | this study |
| | RP | GGTAGATGCGGA TAAAAATGCTTGGTTCGCCCG | |
| F72L | FP | GCATTTTTACCCGCATCTACCTTGAAAATTCCCAATAGCTTGATCG | this study |
| | RP | CGATCAAGCTATTGGGAATTTTCAAGGTAGATGCGGGTAAAAATGC | |
| L158P | FP | GTAGACAGTTTCTGGCCCGACGGTGGTATTCG | this study |
| | RP | CGAATACCACCGTCGGGCCAGAAACTGTCTAC | |
| F156C | FP | GGGCAATGTAGACAGTTGTTGGCTCGACG | this study |
| | RP | CGTCGAGCCAACAACTGTCTACATTGCCC | |
| F156V | FP | GGGCAATGTAGACAGTGTCTGGCTCGACGG | this study |
| | RP | CCGTCGAGCCAGACACTGTCTACATTGCCC | |
| G160C | FP | CAGTTTCTGGCTCGACTGTGGTATTCGAATTTCGG | this study |
| | RP | CCGAAATTCGAATACCACAGTCGAGCCAGAAACTG | |
| G131S-out | FR | AGCGAGGCACGTATGAGCAAG | this study |
| | RP | AATTTGGCGGGCAAATTCTTG | |
| N146S-out | FP | GTGAGGACATTTCGGGCAATG | this study |
| | RP | TACCATAATCGAAAGCATGTAGC | |
| OXA-48-pro-f | | TTGACACAGATATTTATGATATAATAACTGAGTAAGCTTAACATAAGGAGGA AAAACATATGCGTGTATTAGCCTTATCGG | (Jensen & Hammer, 1998) |
| cat-r | | GTAGCACCAGGCGTTTAAGG | this study |
| p15A46 | | TCGTATGGGGCTGACTTCAG | this study |

[a]FP= forward primer, RP reverse primer

**Table S4.** Enzyme concentrations (nM) for steady-state kinetics

| OXA-48 variants | Ampicillin | Piperacillin | Ceftazidime | Cefepime | Imipenem | Meropenem |
|---|---|---|---|---|---|---|
| wild-type | 1 | 1 | 100 | 100 | 25 | 25 |
| L67F | 10 | 100 | 500 | 500 | 50 | 50 |
| P68S | 10 | 100 | 500 | 500 | 50 | 50 |
| F72L | 10 | 100 | 1000 | 1000 | 50 | 50 |
| F156C | 10 | 100 | 500 | 500 | 50 | 100 |
| F156V | 100 | 100 | 500 | 500 | 50 | 100 |
| L158P | 100 | 100 | 500 | 500 | 50 | 50 |
| G160C | 10 | 100 | 500 | 500 | 50 | 50 |
| F72L/G131S | 100 | 100 | 500 | 500 | 50 | 50 |
| N146S/L158P | 100 | 100 | 500 | 500 | 50 | 100 |

**Table S5.** X-ray data collection and refinement statistics for the OXA-48 variant L67F in complex with hydrolysed ceftazidime**.** Values in parenthesis are for the highest resolution shell.

| | OXA-48:L67F |
|---|---|
| PDB no. | 7ASS |
| Diffraction source | BL14.2, Bessy |
| Wavelength (Å), Temperature (°C) | 0.9184, -173 |
| Crystal-detector distance (mm) | 174.9 |
| Rotation range per image (°), total rotation range (°) | 0.10, 190 |
| Space group | $P2_12_12_1$ |
| *a*, *b*, *c* (Å) | 88.56, 108.48, 125.42 |
| Resolution range (Å) | 50.00-1.91 (1.94-1.91) |
| No. of unique reflections | 94247 (4653) |
| Multiplicity | 7.1 (7.3) |
| Completeness (%) | 100 (99.9) |
| $R_{merge}$ (%) | 16.5 (187.0) |
| $R_{pim}$(%) | 9.9 (111.0) |
| Mean $\langle I/\sigma_{(I)} \rangle$ | 9.6 (1.1) |
| C $_{1/2}$ | 0.997 (0.583) |
| Overall *B*-factor from Wilson plot (Å$^2$) | 14.4 |
| Resolution range (Å) | 23.56-1.91 |
| Final $R_{work}$ (%) | 18.9 |
| Final $R_{free}$ (%) | 22.98 |
| Molecules in asymmetric unit | 4 |
| No. of non-H atoms (all protein chains) | 8896 |
| -Ions (Cl) | 5 |
| -Ligand (2 ceftazidime molecules) | 76 |
| -Water | 797 |
| R.m.s. deviations | |
| -Bonds (Å) | 0.011 |
| -Angles (°) | 1.028 |
| Average B-factors (Å$^2$) | 30.3 |
| - Protein chains A/B/C/D | 31.1/26.9/29.4/30.9 |
| -Ion (Cl) | 46.9 |
| -Ligand (2 ceftazidime molecules) | 55.5 |
| -Water | 35.5 |
| Ramachandran plot | |
| Most favored (%) | 96.9 |
| Allowed (%) | 3.1 |

### Supplementary figures

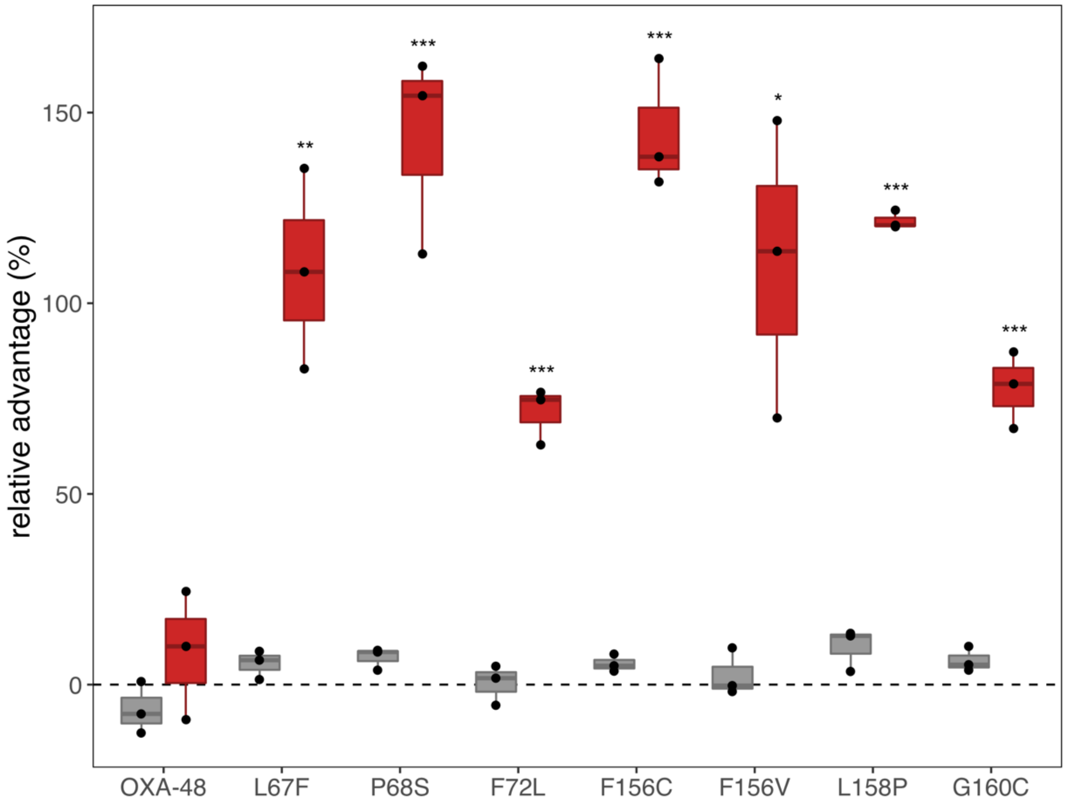


**Figure S1.** Head-to-head competitions, between *E. coli* MG1655 *mal*$^+$ and MG1655 $\Delta$*malF* expressing wild-type and allele variants of OXA-48, conducted without (grey) and at sub-MIC (red) of ceftazidime. While expression without selection pressure was neutral for all alleles, at sub-MIC, all allele variants showed fitness benefits over the wild-type allele. The dots represent biological replicates and significantly different averages, compared to OXA-48 in the presence of ceftazidime (0.06 mg/L), are marked with * (P< 0.05), ** (P<0.01) and *** (P<0.001).

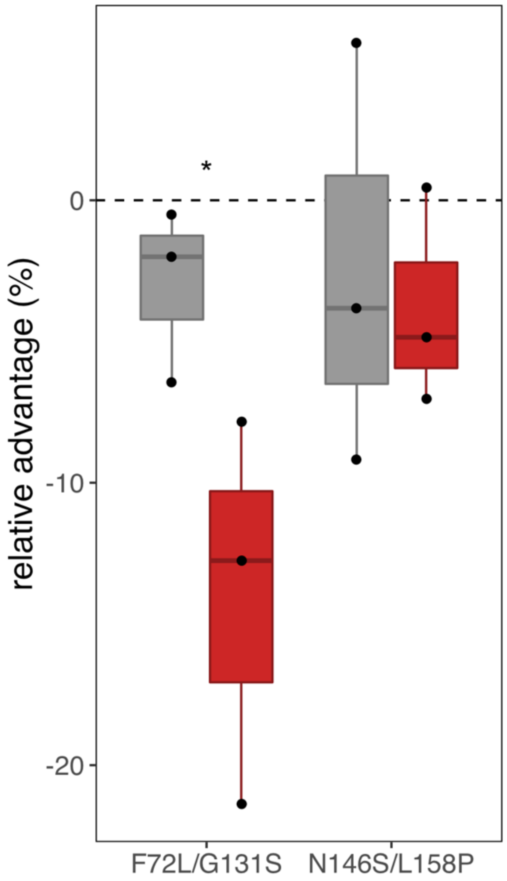


**Figure S2.** Head-to-head competitions between MG1655 expressing F72L versus F72L/G131S and L158P versus N146S/L158P. G131S and N146S did not improve bacterial fitness at sub-MIC ceftazidime. The dots represent biological replicates. Significant differences are indicated as * representing P< 0.05.

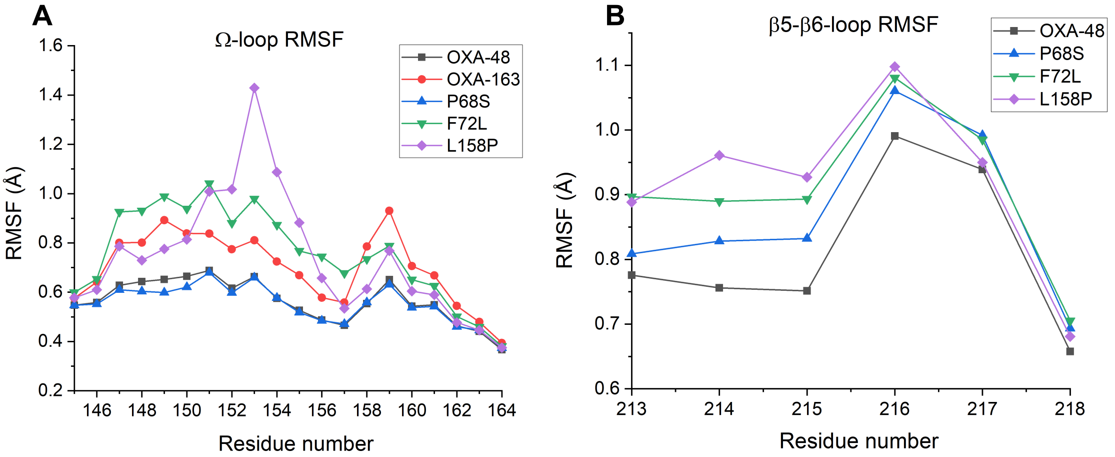


**Figure S3.** Root mean square fluctuations (RMSFs) for the clinical variants OXA-48 and OXA-163 as well as for a sub-set of OXA-48 variants: P68S, F72L and L158P. RMSFs for residues in the Ω-loop (A) and for the β5-β6 loop (B).

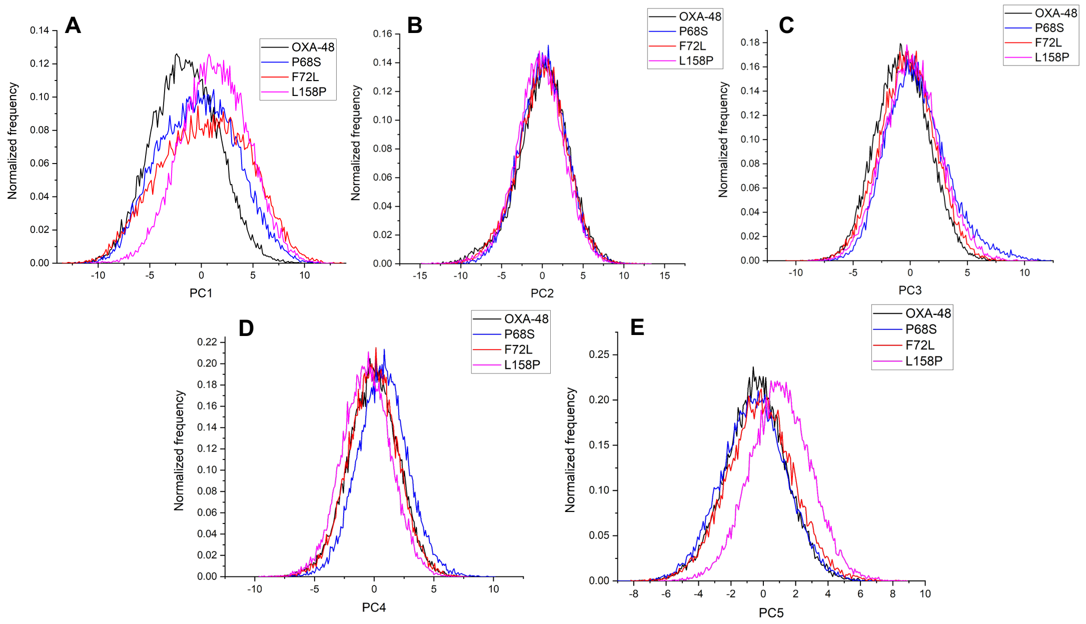


**Figure S4**. Normalized histograms of PC1-PC5 (A to E) for OXA-48 and the variants P68S, F72L and L158P. The histograms are calculated using 200 bins per enzyme.

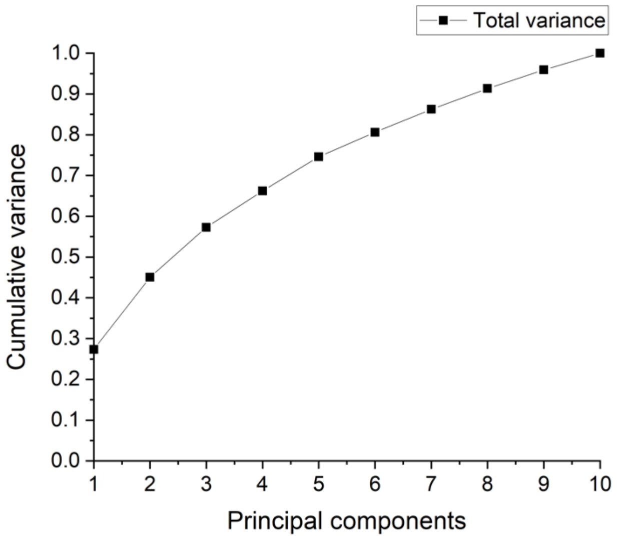


**Figure S5.** Cumulative variance covered by the ten principal components.

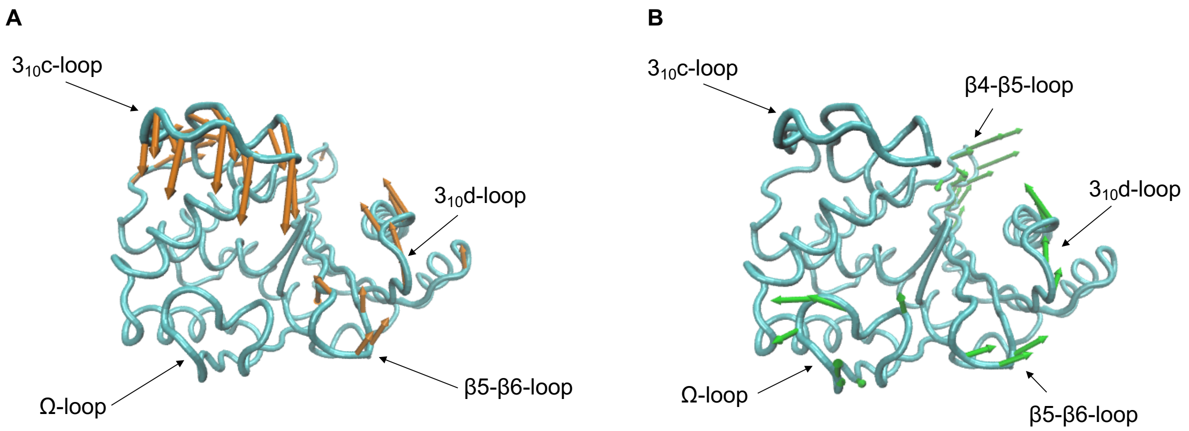


**Figure S6.** Dynamic differences between wild-type OXA-48*,* P68S, F72L and L158P captured by PC1 (A) and PC5 (B). Arrows indicating Cα-movement in PC1 (A) and PC5 (B), arrow direction and size indicating direction of the eigenvector and magnitude of the corresponding eigenvalue (arrows only shown for atoms with eigen values > 2.5 Å).

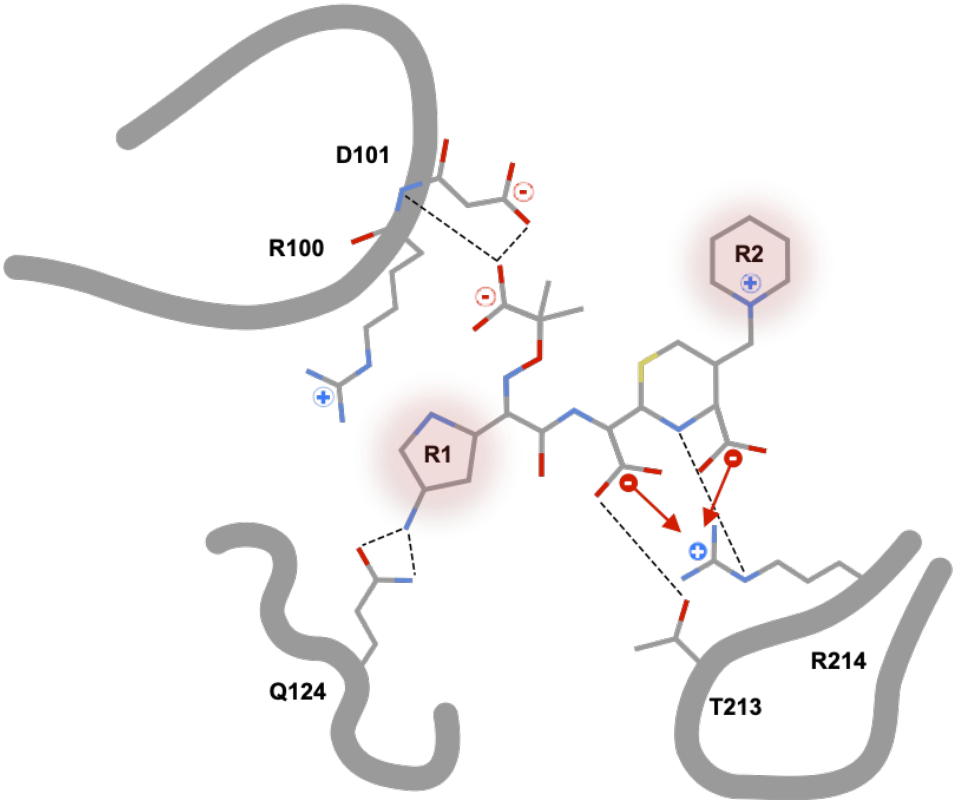


**Figure S7:** Schematic representation of hydrolysed ceftazidime in front of the active site of the OXA-48 variant L67F (based on PDB no. 7ASS). The ceftazidime side chains R1 and R2 are labelled and marked. For R2, no electron density and therefore no interactions were detected. Hydrogen bonds from ceftazidime to D101, Q124, T213 and R214 are represented with dashed lines. The salt bridges to R214 are indicated with arrows.
